# cytoGPNet: Enhancing Clinical Outcome Prediction Accuracy Using Longitudinal Cytometry Data in Small Cohort Studies

**DOI:** 10.1101/2025.05.01.651729

**Authors:** Jingxuan Zhang, Liwen Sun, Neal E. Ready, Wenbo Guo, Lin Lin

## Abstract

Cytometry data, including flow cytometry and mass cytometry, are now standard in numerous immunological studies, such as cancer immunotherapy and vaccine trials. These data enable the monitoring of an individual’s peripheral immune status over time, providing detailed insights into immune cells and their role in clinical outcomes. However, traditional analyses relying on summary statistics, such as cell subset proportions and mean fluorescence intensity, may overlook critical single-cell information. To address this limitation, we introduce cytoGPNet, a novel approach that harnesses extensive cytometry data to predict individual-level outcomes. cytoGPNet is designed to address four key challenges: (1) accommodating varying numbers of cells per sample; (2) analyzing the longitudinal cytometry data to understand temporal relationships; (3) maintaining robustness under the constraints of limited individual samples in immunological studies; and (4) ensuring interpretability to facilitate biomarker identification. We apply cytoGPNet to data from multiple diverse studies, each with unique characteristics. Despite these differences, cytoGPNet consistently outperforms other popular methods in terms of prediction accuracy. Moreover, cytoGPNet provides interpretable results at multiple levels, offering valuable insights. Our findings underscore the effectiveness and versatility of cytoGPNet in enhancing the analysis of cytometry data and its potential to advance immunological research.

## Introduction

Routinely monitoring the immune responses in individuals to diseases or interventions, including therapies and vaccinations, holds paramount significance in clinical trials, treatment modalities, and vaccine development. This imperative involves not only identifying immune responses that confer protection against infection or disease but also predicting an individual’s response to intervention. The evolution of cytometry technologies, including flow cytometry and mass cytometry (CyTOF), empowers researchers to systematically monitor the peripheral immune status of an individual at a reasonable cost. These technologies offer comprehensive insights into immune cell subsets, activation status, polyfunctionality, and other features. With the capacity to measure 10 to 50 parameters (cell markers/variables), encompassing both phenotypic and functional markers, on a large number of single cells (ranging from hundreds of thousands to several millions) for a single sample, such as blood, these multi-dimensional cytometry data provide a rich source of information. This information is invaluable for predicting clinical outcomes, including responses to vaccinations and cancer immune therapies^1–11^.

The conventional analysis for predicting clinical outcomes, employing traditional gating strategies^12^ for cytometry data, can be time-consuming and suboptimal. This approach independently conducts quantitative feature selection and summarization, such as deriving cell subset proportions and mean expression based on gating results, which then serve as a feature vector for outcome prediction. Despite the development of advanced methods to automate manual gating like SPADE, PhenoGraph, FlowSOM, FlowMeans, FlowPeaks, OpenCyto, and others^13–22^, the stepwise approach—deriving summary statistics first and then using them for prediction—may compromise single-cell resolution, potentially introducing variation and obscuring crucial single-cell information relevant to clinical outcomes^23–25^. To address this, approaches that directly utilize single-cell matrix data as input have recently gained popularity as potent tools for predicting outcomes, such as CloudPred^26^, ProtoCell4P^27^, scPheno^28^, DeepGeneX^29^, and ScRAT^30^ for single-cell RNA sequencing data. Notable examples for cytometry data include CellCnn^31^,CyTOF_DL^32^, and CytoSet^33^.

In contrast to the existing applications of statistical and machine learning (ML) methods for cytometry data analysis–such as dimensionality reduction, visualization, identification of cell subsets, and data integration^34,35^– the direct utilization of the single-cell data matrix as input for individual subject outcome prediction presents unique challenges. Firstly, employing advanced ML methods, such as deep neural networks (DNNs), for prediction offers a more flexible model capacity, potentially leading to more accurate outcome prediction. However, training DNNs requires a large number of well-labeled training samples to achieve good prediction performance, posing challenges in studies with a limited number of subjects. Secondly, different samples, represented as single-cell data matrices, often contain varying numbers of cells, hindering the direct use of most existing ML methods without compromising single-cell resolutions. For instance, approaches like summarizing single-cell data by mean marker expressions^29^ or subsampling the cells to achieve an equal number of cells per sample^32,33^ can lead to information loss during cell aggregation and cell resampling from the original data. Thirdly, the black-box nature of DNN classifiers restricts interpretability, hindering the identification of predictive biomarkers. Lastly, and most importantly, existing methods do not directly model the relationships between single-cell data obtained longitudinally, which can be crucial for understanding the effects of diseases or interventions^36–41^. Studies, like immunotherapy trials and early-phase vaccine trials, typically enroll a limited number of individual patients/participants but conduct extensive cytometry-based immune profiling, including longitudinal samples from multiple time points such as pre and post-treatment/vaccination. Consequently, maintaining coherence in the analysis and recognizing the longitudinal relationships between samples is crucial for understanding the effects of diseases or interventions^36,39,41^.

While existing methods address some of the challenges associated with predicting clinical outcomes, they fall short of addressing all the aforementioned obstacles. To bridge this gap, we propose a novel analysis strategy named cytoGPNet, which seamlessly integrate DNNs into the classical Gaussian process (GP) model. By employing cell-level pretraining of an auto-encoder, cytoGPNet learns meaningful cell representations even with small sample sizes (number of subjects). Additionally, the flexibility of the GP model allows for robust handling of cell representations, accommodating variations in cell subsets and the number of cells per sample. Moreover, we incorporate customized attention layers to capture temporal dependency. To further enhance interpretability, we introduce a lightweight prediction model based on GP along with a post-hoc interpretation technique.

## Results

### The cytoGPNet pipeline

As demonstrated in Figure 1, our proposed cytoGPNet architecture has three components: **(1)** a DNN-based auto-encoder (AE) serves as a dimensionality reduction module, encoding single-cell cytometry data into a low-dimensional latent space; **(2)** a Gaussian process (GP) model that can capture correlations between cells within the same individual, across different individuals, and over time, thereby enhancing the model’s ability to capture inter-cell dependencies and temporal relationships for improved prediction outcomes; and **(3)** an attention-based outcome predictor that adaptively summarizes cell-level information from each sample, facilitating the subject’s clinical outcome prediction.

**Figure 1.**
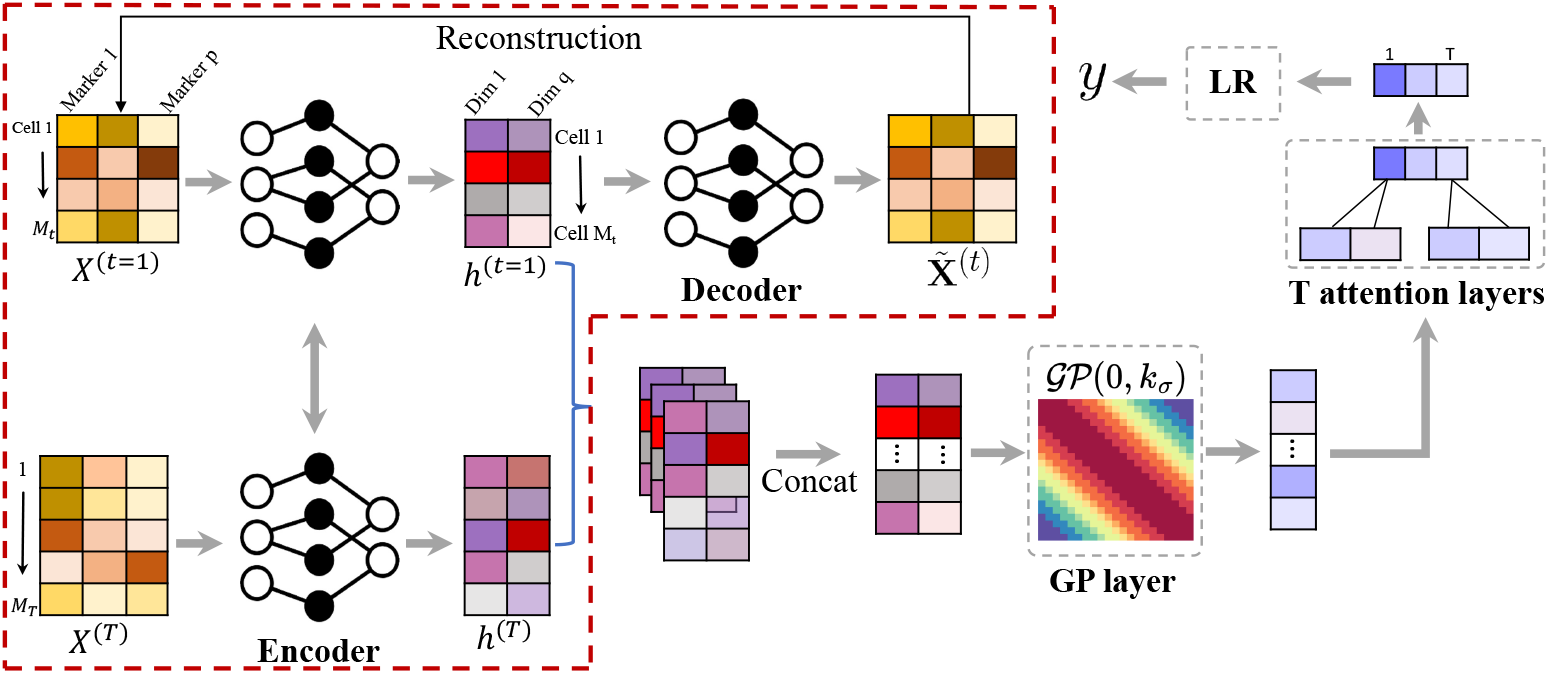
Overview of the proposed cytoGPNet framework. First, we pre-train an auto-encoder model at the single-cell level, using the encoder to obtain a low-dimensional representation for each cell (highlighted by the red dashed line). Next, the encoder output is fed into our GP model, which captures correlations across subjects and time points. We then obtain a sample representation by aggregating the cell representations using multiple attention layers, with each attention layer applied to the cell representations at a specific time point. Finally, the sample representation is used as input to a logistic regression for outcome prediction.

More specifically, the DNN-based AE transforms single-cell cytometry data into a lower-dimensional latent space. In contrast to conventional unsupervised dimensionality reduction methods like UMAP^42^ and t-SNE^43^, our AE module is uniquely designed for end-to-end training with the subsequent GP model. The AE is initially pretrained at the cell level without requiring subject outcomes information, fully leveraging the vast number of cells to learn meaningful cell representations even with a limited number of subjects. However, our analysis reveals that while the AE excels at learning single-cell embeddings, it falls short in capturing multi-cell correlations. This is where the GP becomes more well-suited, as it is better equipped for exploring the immunological variation across cells of different subjects and time points. Modeling such variability is crucial for predicting the subjects’ outcomes^36,39,41,44,45^. However, GP requires less noisy input for optimal performance. By combining the AE with GP, we fully harness the GP’s potential for more informative single-cell learning. This design also benefits from automatically adapting the latent space to align with the GP’s kernel function during training, thereby enhancing the GP’s capability to capture correlations among input cells. In our model, we train the latent space as an Euclidean space, employing the Euclidean distance as the distance metric. In the next step, our GP model takes these latent representations as input and outputs a more compressed representation for each cell that also contains information from other cells. In particular, we choose the squared exponential (SE) kernel^46^ for our GP. This kernel is used to extract cell differences or correlations from the AE-derived latent space, which is an Euclidean space.

Finally, in predicting subject-level outcomes, we employ a set of attention layers to adaptively consider pertinent information from all cells of this subject across all time points. The attention layers are followed by a simple classifier—in this case, logistic regression—to generate prediction outcomes. The attention layers are a key feature of cytoGPNet, allowing the model to accommodate varying numbers of cells across samples by automatically learning optimal weights during training. This enhances model flexibility and reduces the complexity of hyper-parameter tuning required for cell summarization per sample. In contrast to basic summary statistics like mean or sum, which assign predefined and fixed weights to each cell, our approach offers a more adaptive and effective method for predicting subject-level outcomes.

Our proposed model is designed for end-to-end training. It incorporates AE, GP and the attention layers to collectively model, learn, and summarize crucial cell information, thereby enhancing prediction accuracy. Additionally, we develop a customized posterior inference and parameter learning method based on inducing points and variational inference^47^, facilitating efficient model training in a first-order optimization method (e.g., stochastic gradient descent^48^ or Adam^49^). Finally, we develop a post-hoc explanation technique to enhance the interpretability of cytoGPNet.

### cytoGPNet enables accurate outcome prediction

We first benchmarked the prediction performance of cytoGPNet against competing methods on five cytometry datasets—three flow cytometry and two cyTOF—spanning various diseases: COVID-19 (SDY1708), influenza (SDY212), HIV (HEUvsUE), non-small cell lung cancer (TOP1501), and Cytomegalovirus (CMV) infection (CMV). These datasets feature diverse dimensionalities (with the number of subjects *N* ranging from 20 to 308 and the marker dimension *p* ranging from 8 to 49) and different temporal dynamics (with time points *T* =1, 2, 3) (**Datasets**). Additionally, we expanded our analysis by incorporating single-cell RNA sequencing (scRNA-seq) data (SC4) to further evaluate cytoGPNet’s predictive performance across different single-cell data types. The summaries of all six datasets are provided in Supplementary Table 1. CellCnn, CyTOF_DL, Cytoset, AE-based method (AE), logistic regression (LR), and random forest (RF) were used for comparison (**Comparison methods**). For each of the datasets, we applied the 5-fold cross-validation strategy to ensure robust model evaluation and to prevent overfitting. The input to our model cytoGPNet is the preprocessed cytometry data (**Data preprocessing**) from all individual subjects, formatted as matrices with columns representing markers and rows representing the individual cells. The outputs of the model are subject-level information of interest, which are binary outcomes for our six datasets.

**Table 1.** Table summarizing the prediction performance of cytoGPNet and competing methods using AUC and F1-score metrics. Evaluation is conducted on three longitidinal datasets (SDY212, TOP1501 and CMV) using 5-fold cross-validation, comparing three scenarios: full dataset (Full data), 50% reduced cells (50% cells), and 10% randomly masked samples from the second time point (Missing time points). The top performance in each column is highlighted in bold. Values are reported as the mean across 5 folds, with the standard deviation scaled by a factor of 100 in parentheses.

| **SDY212** |  | **AUC** |  |  | **F1** |  |  |
| --- | --- | --- | --- | --- | --- | --- | --- |
| **Method** | Full data | 50% cells | Missing time points |  | Full data | 50% cells | Missing time points |
| cytoGPNet | **78.1(10.3)** | **77.7(8.17)** | **67.6(12.3)** |  | **83.6(6.54)** | **84.9(3.75)** | 74.2(8.81) |
| AE | 68.4(7.67) | 70.0(7.76) | 62.7(12.5) |  | 81.1(3.84) | 81.2(2.97) | 62.6(7.74) |
| CyTOF_DL | 56.58(15.29) | 50.47(11.08) | NA |  | 36.28(21.34) | 32.34(19.97) | NA |
| Cytoset | 65.42(17.42) | 62.78(15.98) | NA |  | 48.41(14.93) | 34.67(20.63) | NA |
| CellCnn | 57.55(15.55) | 62.76(2.93) | NA |  | 28.43(26.14) | 38.89(21.87) | NA |
| LR | 59.3(19.6) | 59.3(19.6) | 50.2(11.9) |  | 77.6(4.97) | 77.6(4.97) | 68.5(2.70) |
| RF | 57.8(7.73) | 56.8(4.81) | 46.9(6.22) |  | 81.2(2.00) | 81.2(2.00) | **77.2(4.93)** |
| **TOP1501** |  | **AUC** |  |  | **F1** |  |  |
| **Method** | Full data | 50% cells | Missing time points |  | Full data | 50% cells | Missing time points |
| cytoGPNet | **85.0(33.5)** | **85.0(33.5)** | **68.0(28.0)** |  | **78.3(15.8)** | **86.7(16.3)** | **60.8(12.7)** |
| AE | 78.0(22.8) | 69.5(19.2) | 60.5(29.4) |  | 64.0(25.1) | 52.8(9.88) | 50(13.4) |
| CyTOF_DL | 57.5(44.72) | 58.5(35.95) | NA |  | 32.38(20.52) | 25.71(25.05) | NA |
| Cytoset | 36.00(29.35) | 46.5(29.98) | NA |  | 22.67(21.43) | 29.05(29.76) | NA |
| CellCnn | 60.00(32.06) | 63.00(24.39) | NA |  | 32.38(20.52) | 32.38(20.52) | NA |
| LR | 65.0(14.1) | 50.5(24.5) | 65.0(14.1) |  | 42.7(7.23) | 40.8(6.87) | 42.7(7.23) |
| RF | 59.5(38.5) | 59.5(38.9) | 55.0(11.2) |  | 66.7(NA) | 66.7(NA) | 38.1(11.0) |
| **CMV** |  | **AUC** |  |  | **F1** |  |  |
| **Method** | Full data | 50% cells | Missing time points |  | Full data | 50% cells | Missing time points |
| cytoGPNet | **90.0(22.4)** | **87.6(19.4)** | **72.3(21.8)** |  | **82.7(16.7)** | **84.6(10.1)** | **69.4(22.9)** |
| AE | 80.0(27.4) | 74.8(22.4) | 64.1(10.2) |  | 79.3(21.7) | 72.0(20.1) | 59.3(10.2) |
| CyTOF_DL | 60.00(45.41) | 60.00(45.41) | NA |  | 69.33(21.27) | 27.33(23.85) | NA |
| Cytoset | 56.67(30.84) | 55.00(37.08) | NA |  | 53.80(32.63) | 57.14(32.99) | NA |
| CellCnn | 65.00(48.73) | 61.67(26.09) | NA |  | 62.48(35.62) | 56.67(36.51) | NA |
| LR | 84.0(18.4) | 84.0(18.4) | 59.2(29.7) |  | 82.0(20.5) | 82.0(20.5) | 48.2(31.4) |
| RF | 85.0(22.4) | 81.2(19.7) | 60.2(21.4) |  | 78.0(17.9) | 79.1(19.1) | 59.7(29.3) |

To evaluate the prediction performance, we use the following four metrics: area under the receiver operating characteristic curve (AUC), F1 score (F1), precision, and recall. The AUC measures the overall performance of a model by evaluating the area under the ROC curve, which plots the True Positive Rate (TPR) against the False Positive Rate (FPR) across various threshold settings. An AUC of 0.5 indicates that the model performs no better than random guessing. The F1 Score is a metric that assesses a model’s accuracy by considering both precision and recall. It is the harmonic mean of precision (the proportion of true positive predictions out of all positive predictions) and recall (the proportion of true positive predictions out of all actual positive instances). The F1 Score aims to balance the trade-off between precision and recall, providing a single measure of a model’s ability to correctly classify instances as positive or negative. For all four metrics, the values range from 0 to 1, with 1 indicating a perfect classifier.

Figure 2 presents a detailed comparison of the prediction performance of cytoGPNet against other methods. The bar plots highlight the AUC, F1, precision, and recall for each method, evaluated on four distinct datasets. Our method, cytoGPNet, consistently outperforms the alternatives, as indicated by the highest AUC and F1 scores, demonstrating its superior predictive accuracy. Notably, cytoGPNet shows significant improvement over the AE method, which is limited by its inability to model cell-cell correlations in single-cell inputs. This suggests that the GP model within cytoGPNet effectively models complex cell-cell interactions, thereby enhancing prediction accuracy and contributing to cytoGPNet’s improved generalizability. The other three deep learning-based methods—CellCnn, Cytoset, and CyTOF_DL—exhibit varying degrees of performance but generally show lower AUC and F1 values compared to cytoGPNet. Sometimes, these methods perform even worse than the traditional LR and RF models. For instance, this is observed in CMV dataset, which has a relatively small sample size (*N* = 20) and a higher dimensionality for the markers (*p* = 39). All three methods exhibit lower values than LR and RF for AUC, suggesting their potential limitations in handling datasets with less information in terms of sample size and dimensionality. This highlights the necessity of more robust methods like cytoGPNet that can effectively manage these challenges and provide more accurate predictions. In addition, a comparison of computational time between cytoGPNet and other deep learning-based methods is provided in Supplementary Figure 1. While our method exhibits a slightly longer running time compared to others, the difference is not significant, indicating comparable performance across all methods.

**Figure 2.**
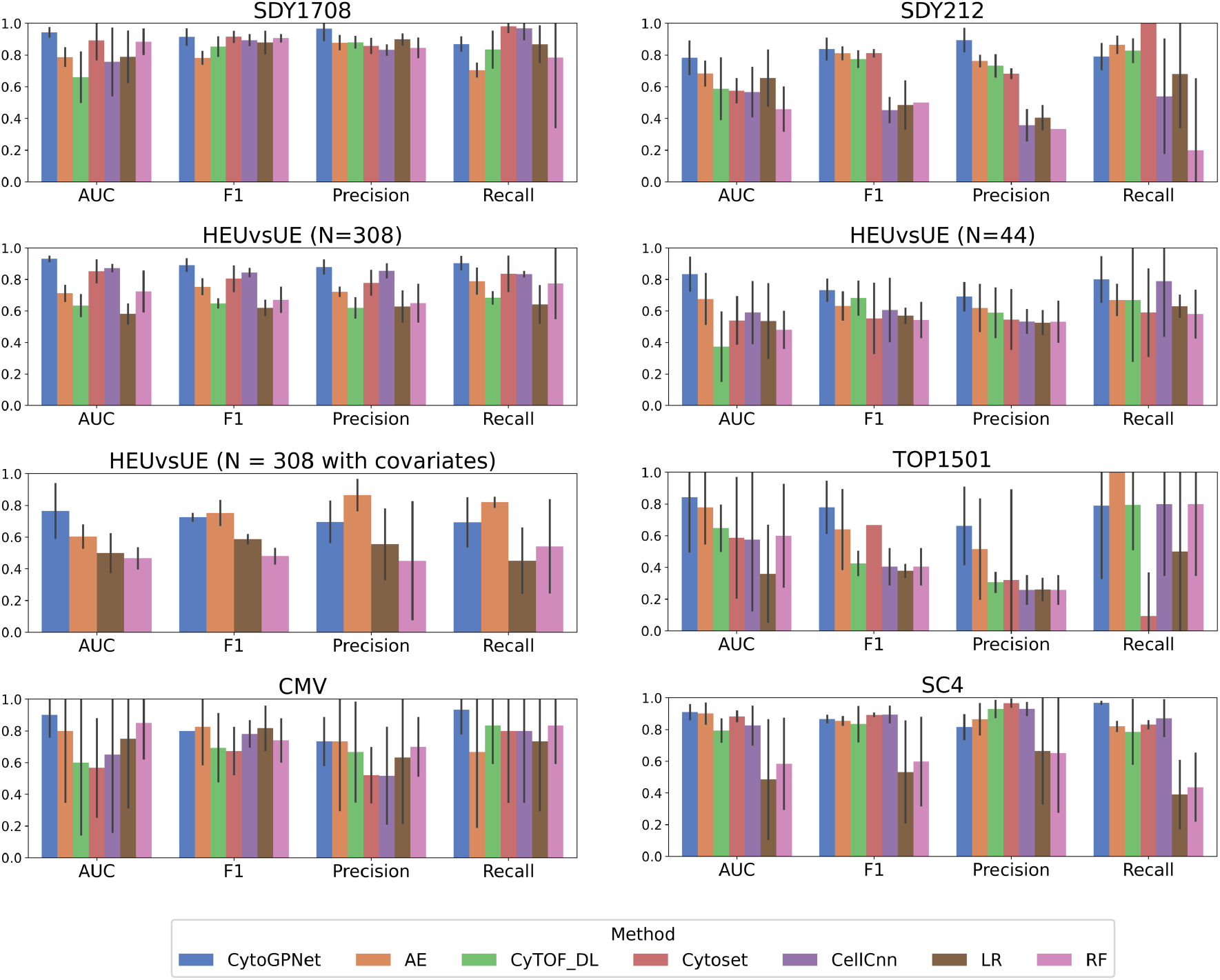
Performance of cytoGPNet and the competing methods measured by AUC, F1, precision, and recall using 5-fold cross-validation on six datasets: SDY1708, SDY212, HEUvsUE, TOP1501, CMV, and SC4. The height of each bar represents the corresponding mean value across the five folds, and the vertical lines (error bars) indicate the standard error.

For HEUvsUE data analysis, we conducted three experiments: (1) aggregating all blood samples across the seven conditions (*N* = 308), (2) using only the unstimulated blood samples (*N* =44), and (3) incorporating condition information for all 308 samples as additional features (*N* =308 with covariates). In the third experiment, we introduced six dummy variables (*I*_1_, *I*_2_, …, *I*_6_) in the final classification layer as covariates in the logistic regression model to account for the seven experimental conditions, with the unstimulated condition serving as the reference group. Each dummy variable corresponds to one of the six stimulation conditions, taking a value of 1 if the sample belongs to that condition and 0 otherwise. This encoding allows the model to estimate the effect of each stimulation condition relative to the unstimulated baseline. The results, shown in the second row and the left panel of the third row of Figure 2, demonstrate that our method remains robust to variations in sample size. However, in the third experiment (*N* =308 with covariates), we observe a slight decrease in the predictive performance of cytoGPNet. This decline is likely due to the lack of significant associations between the six dummy variables and the outcome variable when fitting cytoGPNet to the entire dataset (the smallest p value is 0.24, and the largest is 0.76). Since these covariates do not provide informative contributions to the outcome, their inclusion may introduce unnecessary complexity, ultimately reducing predictive power. Additionally, it is worth noting that none of the benchmarked deep learning-based models can incorporate condition information as covariates into the analysis without modifying the existing code. These findings highlight our algorithm’s adaptability to different data configurations while maintaining its predictive capability.

For scRNA-seq data (SC4), there are two key differences compared to mass and flow cytometry data: they contain significantly fewer cells per sample and exhibit much higher dimensionality due to the large number of genes measured per cell. To accommodate this high-dimensional nature, we modified our AE architecture used for cytometry data by adding three additional fully connected hidden layers with output sizes of 256, 64, and 16. We used the ReLU activation function and trained the autoencoders using the Adam optimization algorithm. The widths of the three hidden layers were pre-specified without additional tuning, informed by prior experience with scRNA-seq data analysis using autoencoders^50^. This was the only structural adjustment made to cytoGPNet, underscoring its flexibility in handling diverse data modalities while maintaining strong predictive performance. To showcase cytoGPNet’s flexibility in handling non-binary outcome data, we further categorized patient outcomes into three groups: healthy, mild or moderate, and severe. To accommodate this multiclass prediction, we extended our model’s final classification layer to a multinomial framework, replacing the sigmoid activation function with a softmax function. This modification preserves cytoGPNet’s core structure while enhancing its adaptability to diverse outcome types. The prediction results, illustrated in Supplementary Figure 2, are presented in the form of a confusion matrix.

We also conducted analyses using cell type proportions as features—after applying a centered log-ratio transformation—to perform prediction via logistic regression with a Lasso penalty. Specifically, for datasets lacking manual gating results (i.e., all datasets except TOP1501), we performed automatic clustering using FlowSOM with the default setting of 100 clusters, allowing the algorithm to capture granular cellular subpopulations. FlowSOM then aggregates these 100 clusters into 10 meta clusters, facilitating higher-level biological interpretation. For longitudinal datasets, clustering was performed separately at each time point, and the proportions of each cluster across all time points were included as covariates in the Lasso regression model. As illustrated in Supplementary Figure 3, the performance of proportion-based prediction varies considerably across datasets, whereas cytoGPNet demonstrates consistent performance. Supplementary Figure 4 further examines the TOP1501 dataset: panel (a) displays p values from univariate analyses—where each cell type proportion is added as a covariate individually—and reveals that none of the gated cell types show a statistically significant association with the outcome. Panel (b) compares the performance of cytoGPNet with predictions based on cell type proportions obtained from FlowSOM (using both 100 clusters and 10 metaclusters) as well as manual gating results. For this dataset, we observe a dramatic decrease in prediction performance when relying solely on cell-type proportion-based analysis. This suggests that while cell type proportions are a commonly used summary statistics for single-cell data, other features, such as marker expression levels, can also provide valuable insights.

**Figure 3.**
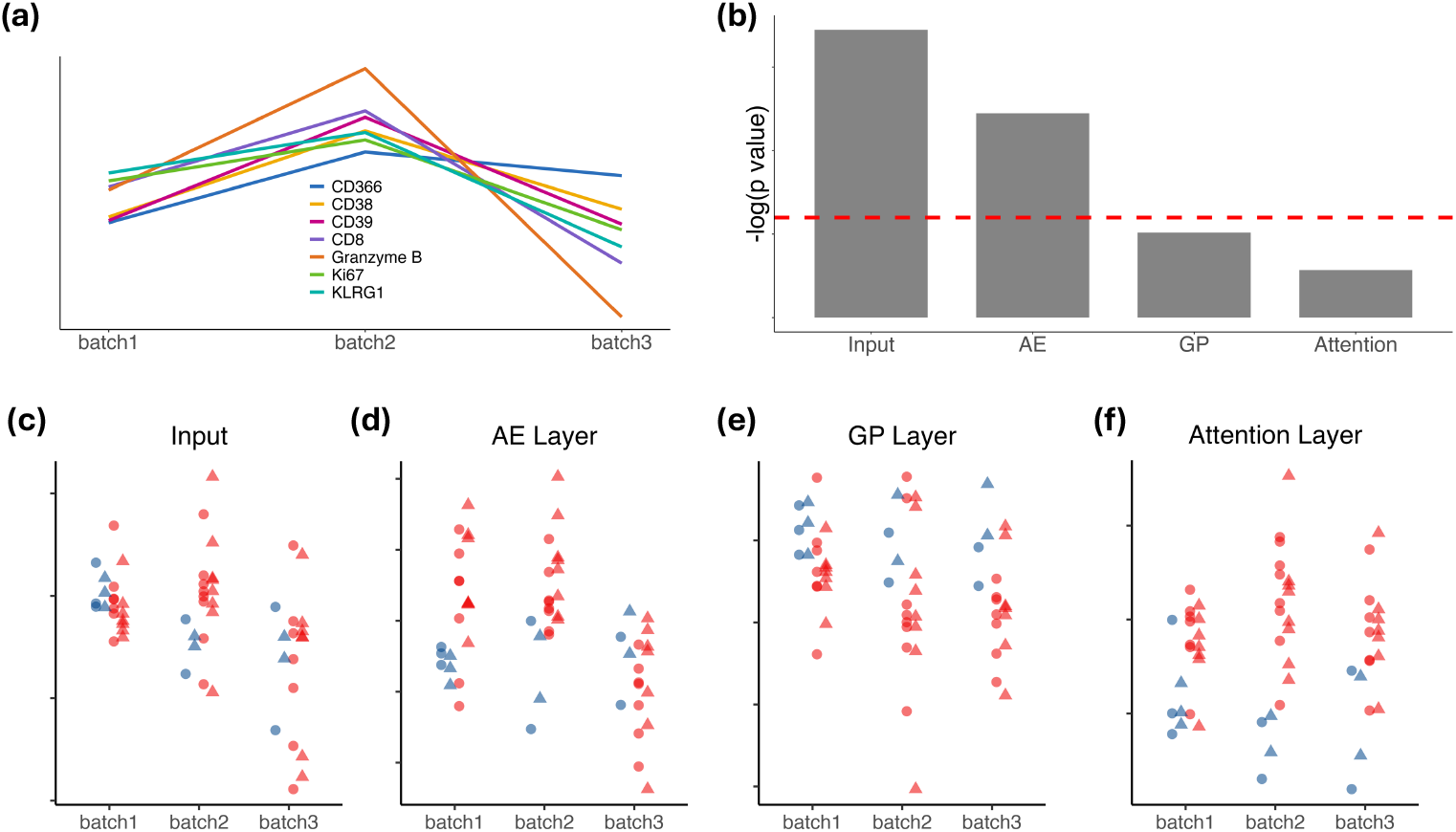
(a) Line plots showing the mean expression of seven selected markers (out of 25) from samples of the same healthy control subject across three batches. (b) Bar plot illustrating the batch effects across different layers in cytoGPNet, measured by the negative logarithm of p values obtained using the Kruskal-Wallis test. The horizontal dotted red line indicates the negative logarithm of 0.05. (c)-(f) Dot plots displaying the activation scores in different layers of cytoGPNet, with each dot representing a sample within a batch, color-coded by the patient’s outcome (blue: responder; red: non-responder), with shapes indicating the time point when the sample was collected (circle: baseline; triangle: post-treatment).

**Figure 4.**
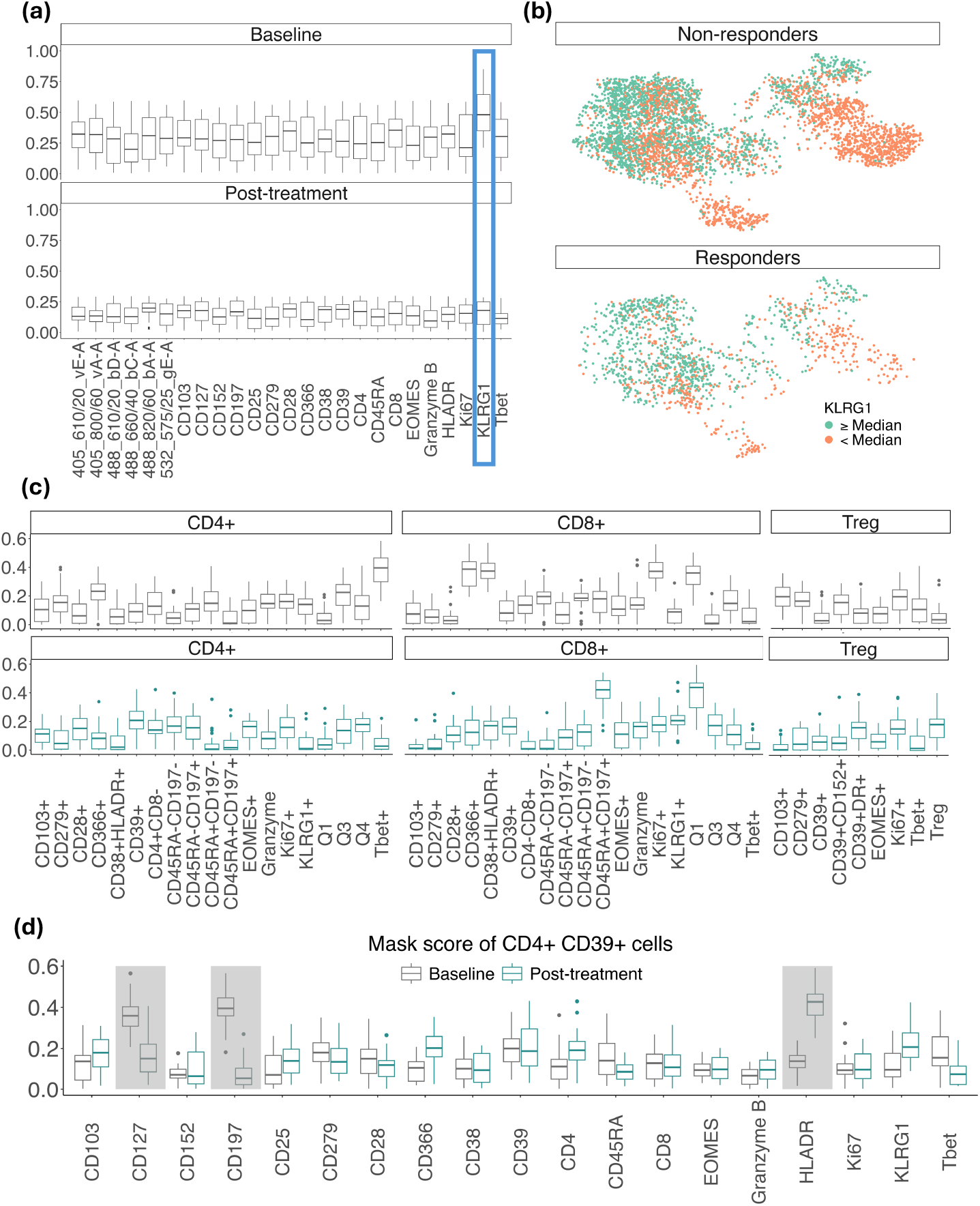
(a) Boxplots visualizing the distribution of masking scores (ranging from 0 to 1) for each marker (x-axis) for baseline data (left) and post-treatment data (right). Marker with the highest masking score is highlighted with blue rectangle. (b) tSNE plots of randomly subsampled cells from baseline samples, shown separately for responders and non-responders and color-coded by KLRG1 expression levels. Cells with KLRG1 expression values above the median are marked in orange, while those below the median are marked in green. (c) Boxplots visualizing the distribution of masking scores for KLRG1 across all cell subsets for both baseline and post-treatment data. (d) Boxplots depicting the distribution of masking scores for CD4+ CD39+ cells.

For both TOP1501 and CMV datasets, which include temporal information and have extremely small sample sizes, all alternative methods experience a decline in terms of AUC. This underscores the critical importance of explicitly modeling longitudinal data to effectively capture temporal dynamics and improve predictive accuracy. We also explored refining our model by explicitly incorporating the longitudinal structure into the additive GP layer. Specifically, we introduced a separate GP designed to learn time-step correlations by obtaining a unified representation of all cells at the same time step and applying a GP to these unified representations. This approach allows the covariance matrix to directly capture temporal dependencies across time points. However, our empirical results did not show a significant or consistent improvement across longitudinal datasets. For instance, while the AUC across five folds increased marginally by 0.2% for the TOP1501 dataset, it declined by 1.4% for the SDY212 dataset and 5% for the CMV dataset. These results indicate that incorporating this additional GP does not improve prediction performance, at least for small sample sizes, suggesting that our original GP formulation is already effectively capturing temporal correlations at a fine granularity. Overall, cytoGPNet’s ability to maintain high prediction performance across various datasets underscores its robustness and adaptability.

### cytoGPNet demonstrates robustness to missing time points and smaller cell numbers

We next evaluated the robustness of cytoGPNet when the data contains missing time points or a reduced number of cells. For this purpose, we utilized the SDY212 and TOP1501 datasets, which both feature cell measurements across two different time points. To simulate a smaller number of cells, we randomly selected 50% of all cells for each blood sample. We then re-trained our model using the same pipeline and assessed its performance using AUC and F1 scores, presented in Table 1. Remarkably, cytoGPNet maintained similar performance on both the full datasets and the datasets with only 50% of the cells. This demonstrates that reducing the number of cells does not significantly impact the accuracy of our model. In contrast, all competing methods either exhibit a decline in performance or continue to perform poorly even with the full datasets.

To evaluate the robustness of the models in the presence of missing time points, we randomly masked 10% of the samples’ data from the second time point, simulating real-world scenarios where patients miss their follow-up visits. Given that CellCnn, CyTOF_DL, and Cytoset cannot handle missing temporal data, we excluded these models from the analysis and focused on comparing cytoGPNet with the remaining methods. More specifically, during the five-fold cross-validation process, we randomly masked 10% of the training samples’ data from the second time point. For subjects with missing cell data, we addressed this by padding with zeros. As shown in Table 1, our model demonstrates superior performance compared to other models, even when accounting for missing time points. Notably, cytoGPNet retains relatively high AUC and F1 values. This indicates the robustness and reliability of our model in handling incomplete datasets while preserving predictive accuracy. This robustness is particularly crucial in longitudinal studies where patient follow-up can be inconsistent, ensuring that cytoGPNet can provide reliable insights even with partial data.

The ability of cytoGPNet to handle missing data and reduced cell numbers without a significant loss in performance indicates its potential applicability in diverse clinical and research settings. For example, in scenarios such as early-phase clinical trials or studies involving rare diseases, where data may be scarce or incomplete, cytoGPNet’s resilience ensures that valuable predictive insights can still be obtained.

### cytoGPNet is able to mitigate batch effect

To evaluate our model’s resilience to batch effects, we utilized the TOP1501 dataset, which consists of samples from three distinct batches in wet lab experiments. Since each batch contained blood samples from the same healthy control subject, we generated line plots for each measured marker by averaging expression levels across cells within the same blood sample. The averaged marker expression was then standardized across the three batches for visualization and comparison. Figure 3(a) illustrates the presence of batch effects. To further assess these effects, we performed the Kruskal-Wallis test, a non-parametric method for comparing multiple groups. This analysis determined whether statistically significant differences existed between sample distributions from different batches. The results indicated that all markers had significant p values well below 0.05, confirming the presence of batch effects across the measured markers.

Next, following the methodology outlined in^32^, We first derived the activation score for each layer of the model. For a given layer, the score is calculated by averaging the output values for each cell after it passes through that layer. This average is computed across all cells and output dimensions within each sample. This score provides a scalar summary measure of the model’s internal representation per sample at different stages of cytoGPNet. Thus, we assessed cross-batch heterogeneity in each layer of cytoGPNet using the Kruskal-Wallis test. Our results, shown in Figure 3(b), indicate that heterogeneity gradually decreases across the layers: it is strongest at the input layer but becomes insignificant in both the GP and attention layers. This trend is further visualized in Figure 3(c)-(f). Each dot represents a single flow cytometry sample, color-coded by the patient’s outcome, with shapes indicating the time point when the sample was collected. These plots visualize how samples from different batches are distributed across cytoGPNet ‘s layers, highlighting how the model processes and integrates information. Starting at the input layer (Figure 3(c)), the blue (responders) and red (non-responders) dots are intermixed both within each batch and cross batches, showing no discernible pattern or separation. However, as cytoGPNet progresses through the subsequent layers, a gradual trend emerges. The blue and red dots begin to separate more distinctly within each batch. For example, in the GP layer, the blue dots tend to have higher values than the majority of the red dots across all three batches, with this separation becoming more pronounced compared to the input and AE layers. This suggests an increasing ability of cytoGPNet to distinguish between responders and non-responders as the data is processed through its layers. By the attention layer (Figure 3(f)), the separation between the blue and red dots becomes even more distinct. For instance, in batch 2, the distinction between the two groups is noticeably clearer than in the preceding GP layer, indicating that the model has effectively captured and utilized relevant signals to differentiate patient outcomes. This progression underscores the model’s learning progression. This trend suggests that cytoGPNet effectively mitigates the batch effects as the data progresses through the network layers. Despite the presence of batch effects, cytoGPNet reliably predicted clinical outcomes at the individual subject level, highlighting its ability to discern meaningful signals amidst batch-related noise.

### cytoGPNet suggests potential biomarker dynamics

Identifying blood-based biomarkers to direct immune therapy for cancers remains a major area of unmet need. In non-small cell lung cancer in particular, obtaining adequate tumor biopsies for immune profiling in a safe and timely manner is often not feasible. Traditional immune profiling of blood has limitations for identifying immune cell subsets associated with benefits from programmed death 1 (PD-1) checkpoint therapy. To uncover the biomarkers associated with cytoGPNet’s high predictive accuracy on the TOP1501 dataset, we employed our model’s masking (explanation) algorithm. This algorithm was applied iteratively to gain insights into the potential dynamics of the biomarkers: first on baseline data and then on post-treatment data. The results, as depicted in Figure 4(a), reveal that Killer cell lectin-like receptor G1 (KLRG1) emerges as the most predictive biomarker for baseline data. Figure 4(b) presents tSNE plots of randomly subsampled cells from baseline samples, shown separately for responders and non-responders, color-coded by KLRG1 expression levels. Cells with expression values above the median are marked in orange, while those below the median are marked in green. Notably, at least two distinct clusters of cells (in the bottom left and bottom right regions of the figure) are observed. In these clusters, non-responders exhibit a higher proportion of cells with elevated KLRG1 expression compared to responders, suggesting a potential association between KLRG1 expression and response status. Interestingly, none of the markers individually stand out as highly predictive for post-treatment data alone. As a validation, we have also included six markers (the first six markers in Figure 4(a)) in our analysis, which were not used in the gating process for identifying T cells. As expected, our analysis did not identify these six markers as predictive biomarkers, suggesting that the markers selected by our model are relevant to predictive power. Among these, Violet E and A (`405_610/20_vE-A`, `405_800/60_vA-A` ) were part of the Innate panel but not the T cell panel, while Blue A, C, D, and Green E (`488_610/20_bD-A`, `488_660/40_bC-A`, `488_820/60_bA-A`, `532_575/25_gE-A`) were not used in either panel and served as empty channels. These empty channels were included in the analysis as noise without compensation. Additionally, we performed a separate analysis excluding all six markers, yielding a mean AUC of 86.19 (SD = 0.29) and a mean F1-score of 81.74 (SD = 0.157) across 5-fold cross-validation—results unchanged from the analysis including these markers. This demonstrates that cytoGPNet effectively identifies key predictive markers.

Our finding in KLRG1 is consistent with existing research findings, such as documented in published study^51^. KLRG1, identified as a co-inhibitory receptor for natural killer (NK) cells and antigen-experienced T cells, has been implicated in immune regulation in patients with non-small cell lung cancer. Studies have shown that KLRG1 knockdown affects tumor cell proliferation, highlighting its potential as a therapeutic target. While we found no predictive markers post-treatment, this does not necessarily imply the absence of differential marker expression between responder and non-responder groups post-treatment. It may suggest that post-treatment differences may be subtler compared to baseline. One possible reason for this could be the administration of 200 mg intravenous pembrolizumab over two cycles. Pembrolizumab functions by blocking interactions between PD-1 on T cells and its ligands PD-L1 and PD-L2 on tumor cells, thereby restoring effective T cell responses against cancer^52^. This mechanism could lead to a reduction in individual-level differences post-treatment, resulting in a more uniform count of effective T cells across patients and consequently less variability in expression data among all patients. Our subsequent analysis further supports this interpretation.

We next applied our model’s explanation algorithm to different cell subsets identified by manual gating, leading to the identification of multiple cell subset-specific predictive biomarkers. Among them, figure 4(c) top panel shows five cell subsets where KLRG1 emerges as the most predictive biomarker for baseline data. Interestingly, the bottom panel of Figure 4(c) demonstrates that while KLRG1 is not a predictive biomarker for post-treatment data when considering all cells collectively, it remains predictive for patient outcomes for two specific cell subsets: CD8+ CD45RA+ CD197+ and CD8+ CD38-HLADR+ cells (annotated as CD8+ Q1 in the figure).

For CD8+ CD45RA+CD197+ cells, the biological function has been well explained in mice data. KLRG1, defined as MPEC in mice, is known to be highly expressed in noncytotoxic cells with memory functions. These cells possess high levels of CD45RA and CD197 and serve as markers for distinguishing cytotoxic cells within the CD8 population^53^. Noncytotoxic cells, also known as “natural killers”, play crucial roles in protective immunity against a wide range of pathogens and tumors^54,55^. In contrast, cytotoxic cells are more effective in eliminating cancer cells as tumor antagonist effectors^56^. Therefore, responders are expected to exhibit lower proliferation of KLRG1 expression in these cells to maintain tumor suppression, especially after pembrolizumab treatment, as supported by our data (not shown in the figure). This finding aligns with a novel treatment approach that combines the anti-cytotoxic T-lymphocyte–associated antigen-4 monoclonal antibody quavonlimab with pembrolizumab to enhance safety and efficacy in treating extensive-stage lung cancer^57^. For CD8+ CD38-HLADR+ cells, KLRG1 appears to be a predictive biomarkers for both baseline and post-treatment data. This observation aligns with previous studies that the percentage of PD-1+ CD8 T cells that were KLRG1-negative was strongly associated with response in both pre-treatment and 2-week on-treatment blood samples^58^. Given that PD-1 positive CD8 T cells are significantly enriched in the specified gating (CD38+ HLADR+), it is plausible to infer that KLRG1 can be a critical marker of terminal differentiation^58,59^.

Figure 4(d) reveals that within the CD4+ CD39+ cell subset, both CD127 and CD197 markers are particularly predictive of outcomes at baseline compared to other markers. Previous research has identified CD127 as a crucial factor in the dynamic regulation of the T-cell compartment, playing a key role in the maintenance of memory T cells^60^. CD197, on the other hand, is predominantly expressed in naive and memory cells, highlighting its importance in these populations^61^. Furthermore, for post-treatment data, HLADR emerges as the most predictive marker within the CD4+ CD39+ cell subset. HLADR is a significant marker involved in antigen presentation to T-cell receptors (TCRs) on T-helper cells, which subsequently leads to antibody production^62^.

We also compared our findings with other competing methods. LR fails to detect differentially expressed markers for prediction using a z-test on the parameters. Supplementary Figure 4(c) illustrates the variable importance of RF for both baseline and post-treatment samples. CD197 emerges as the most important marker for baseline data. However, another marker, 448_660/40_bC*−* A, also appears highly important, despite not being used in the gating process for identifying T cells. Additionally, the relatively low prediction accuracy for RF further undermines the reliability of its importance computation. CytoSet lacks interpretability features, making it challenging to discern the significance of individual markers. In contrast, CellCnn provides cell type-specific filter responses, allowing for differentiation of outcomes based on variations in cell filter response values between outcome groups. Supplementary Figure 4(d) shows the boxplots of cell filter response values for each cell subset, according to manual gating, for both baseline and post-treatment data. None of the cell subsets show a significant difference between responder and non-responder groups. CyTOF_DL uses a permutation-based method to interpret the model, involving up-sampling of cells and a decision tree algorithm to select significant markers for each cell subset. However, due to its high computational demands, we were unable to obtain any results after 24 hours of computation. We also compared our findings with diffcyt, an R package designed for the differential analysis of cytometry data^63^. It provides a framework for detecting differentially abundant cell populations and differentially expressed markers across experimental conditions. It outputs p values for each marker across all cell types, indicating whether a marker is differentially expressed within a specific cell type. Additionally, the Benjamini-Hochberg (BH) procedure is applied to adjust for multiple testing. Supplementary Figure 5 presents a heatmap of adjusted p values from the differential abundance analysis across manually gated cell types and markers under baseline and post-treatment conditions. Overall, both diffcyt and our approach indicate that differential patterns are more pronounced in the baseline data than in the post-treatment data. Furthermore, both methods identify KLRG1 as a significant marker, consistently exhibiting differential expression across most gating labels. Notably, diffcyt detects a larger number of differentially expressed markers, which may be due to its differential analysis within manually gated, overlapping subsets of cells that are not mutually exclusive. In this context, multiple testing correction methods such as the BH procedure tend to be less conservative than when p-values are independent^64–67^. This likely contributes to diffcyt identifying more differentially expressed markers compared to our analysis method. Furthermore, the significant adjusted p values are generally close to 0.05, suggesting that applying a more conservative adjustment method could further reduce the number of differentially expressed markers.

Lastly, we also applied cytoGPNet to the analysis of CMV data. CMV infection significantly impacts immune system dynamics in renal transplant recipients, primarily through the activation and expansion of cytotoxic lymphocytes. A study by Ishiyama et al.^68^ showed that CMV-infected patients exhibit increased expression of proliferation and cytotoxicity markers, such as Ki67, Granzyme B, NKG2C, and CD57, largely due to the adaptive response of natural killer (NK) cells and CD8+ T cells in controlling viral replication. Interestingly, similar expression trends were observed in non-CMV-infected renal transplant patients, suggesting that immune activation may be influenced by additional factors beyond CMV infection. In our analysis of the CMV dataset as shown in Supplementary Figure 6, we identified NKG2C and CD57 as highly predictive markers for CMV classification at day 0, highlighting their potential role in early immune responses to CMV infection. Furthermore, Ki67 was found to be a predictive marker in the post-viremia stage, suggesting its involvement in the proliferative response of cytotoxic lymphocytes following viral clearance. However, contrary to the original study, which identified *FcεRIγ* as a differentially expressed marker in CMV-infected patients, our analysis did not replicate this finding. This discrepancy may stem from differences in patient cohorts and underlying biological variations, which merit further exploration.

## Methods

### The cytoGPNet model

#### Model design

Formally, consider a study comprising *N* individual subjects. Each subject has immune cells measured at time points *t* =1, …, *T* . For instance, with *T* =2, *t* =1 could correspond to the baseline measurement, and *t* =2 corresponds to the measurement after the intervention. Thus, we denote a dataset 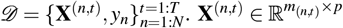 is the single-cell cytometry data for subject *n* measured at time point *t*, where *m*_(*n*,*t*)_ is the number of cells (rows) for **X**^(*n*,*t*)^, and *p* is the number of measured markers for each cell. *m*_(*n*,*t*)_ typically varies across subjects and time. We define 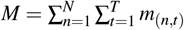 as the total number of cells for the entire dataset *𝒟. y*_*n*_ denotes the outcome for patient *n*. For example, *y*_*n*_ can be a binary value with 1 (responder/protected) or 0 (non-responder/non-protected) in an immunotherapy study/vaccine trial setting. We let 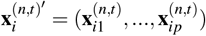 be the *i*th row (cell) of **X**^(*n*,*t*)^, where ^*′*^ denotes transpose. We let **X** denote all cells in the dataset *𝒟*.

As demonstrated in Figure 1, our proposed model combines DNNs with the classical GP model. Our rationale is: (1) Non-parametric models are often preferred in longitudinal study designs because they require fewer assumptions about the underlying mechanism that generates the data. GP is a principled framework for learning non-parametric models probabilistically^69–71^. Thus, by utilizing all given single-cell data as input, GP learns a unified covariance matrix that captures the correlations between cells within the same subject, across subjects and time, thereby improving the modeling of cell dependency and temporal relationship and the final prediction result; (2) The property of GP that any subset of random variables follows a joint Gaussian distribution, regardless of the number and order of variables in the set, enables the construction of a robust model that can handle variations in the number of cells per sample. This feature is particularly advantageous in studying immune responses where cell counts and cell heterogeneity vary across samples. (3) To further strengthen the capabilities of GP, we aim to enhance the robustness and informativeness of single-cell representations by combining GP with the AE. By training AE jointly with the GP model, the AE can learn an adaptable latent space optimized for the subsequent GP model. In other words, the AE model will be trained to convert noisy input into a latent space that aligns with a Gaussian distribution, making it appropriate for GP inputs. This property ensures that the model remains effective even if the original input does not follow a Gaussian distribution. Furthermore, GP and AE together transform the original input of each cell from a vector into a scalar representation, effectively reducing the dimensionality and making the problem easier for the subsequent outcome prediction model.

Specifically, each cell *i* from subject *n* and time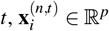, is first fed into an AE model consisting of an encoder *g*(*·*) and a decoder *h*(*·*). The encoder is a multi-layer perception (MLP) that transforms 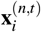 into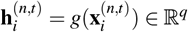, where *q < p*. As shown in Figure 1, the decoder has a symmetric structure as the encoder, which reconstructs the input using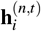, i.e.,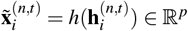. The AE is first pre-trained by minimizing the reconstruction error, i.e, 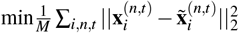. Then, we preserve only the encoder and apply our customized GP model to the encoder outputs **H**. Here, we use **H** to represent the encoder outputs for all cells in the dataset, where 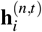 is the representation for cell 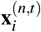.Formally, GP defines a prior of a non-parametric function *f ∼ 𝒢 𝒫* (0, *k*_*σ*_)^1^, where *k*_*σ*_(*·, ·*) is a positive semi-definite kernel function. Here, we use the SE kernel function with *σ* as the scale parameter, and we set the variance to 1 by default. Any finite collections of **f** *∈* ℝ ^*M*^ follows a multivariate Gaussian distribution (**f** | **H**) *∼ N*(**0**, *K*_*HH*_). Here, *K*_*HH*_ *∈* ℝ ^*M×M*^ is the covariance matrix with (*K*_*HH*_)_*ij*_ =*k*_*γ*_ (**h**_*i*_, **h** _*j*_). For simplicity, we omit (*n, t*) and use **h**_*i*_, **h** _*j*_ to represent the representation for any two cells from **H**.

So far, we have obtained a compressed representation for each input cell. To enable prediction at the subject level, we first need to derive a subject-level representation from the output of our GP model. Instead of using simple summary statistics, we utilized multiple attention layers to obtain a representation for each subject. The attention layer can learn to automatically adjust the weight for each cell in the subject representation and their contribution to the final outcome prediction. It enables unique cell weights for different subjects and avoids manual effort in determining the proper cell weights.

Specifically, given the GP outputs of all cells in each subject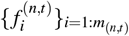, where 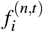 is a scalar, we first split it into *T* sub-vectors according to the temporal information. In other words, the representations of cells from the same subject at the same time are concatenated as the sub-vector. Then, we input each sub-vector into an individual attention layer and output a scalar presentation, denoted as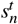. Finally, we concatenate the attention outputs into a subject representation 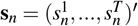.The rationale for having individual attention for each time is to fuse the cell information at the same time and help the subsequent classification model to better capture the correlations across times. Finally, we let *y*_*n*_|**s**_*n*_ follow a Bernoulli distribution with 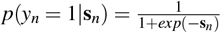.

#### Learning and inference

Direct inference of our model requires *𝒪*(*M*^3^) computational complexity. To improve scalability, we adopt the inducing points method^47^. This method simplifies the posterior computation by reducing the effective number of samples in **X** from *M* to *O*, where *O << M* is the number of inducing points. More specifically, we define *O* inducing points, where each inducing point is **z**_*i*_ *∈ R*^*q*^, and we let **u**_*i*_ be the GP output of **z**_*i*_. Then, the joint prior of the GP outputs **f** on actual inputs **H** and the outputs **u** for inducing points **Z** and the conditional prior **f**|**u** are given by:

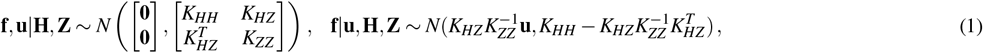

where *K*_*HH*_, *K*_*HZ*_, *K*_*ZZ*_ are the covariance matrices. With inducing points, we only need to compute the inverse of *K*_*ZZ*_, which significantly reduces the computational cost from *𝒪*(*M*^3^) to *𝒪*(*O*^3^). For simplicity, we omit the *n, t* from the GP input **H** and output **f**.

So far, our model has introduced the following parameters: AE parameters Θ_*n*_, GP parameters Θ_*k*_, attention layer and prediction model parameters Θ_*p*_, and inducing points **Z** =*{* **z**_*i*_*}* _*i*=1:*O*_. To learn these parameters, we follow the idea of empirical Bayes^70^ and maximize the log marginal likelihood log *p*(*y*| **X, Z**, Θ_*n*_, Θ_*k*_, Θ_*p*_). Maximizing this log marginal likelihood is computationally expensive and, more importantly, intractable for models with non-Gaussian likelihood. To provide a factorized approximation to marginal likelihood and enable efficient learning, we assume a variational posterior over the inducing variable *q*(**u**) *∼ N*(*µ*, Σ) and a factorized joint posterior *q*(**f, u**) =*q*(**u**)*p*(**f**|**u**), where *p*(**f**|**u**) is the conditional prior in Eqn. (1). By Jensen’s inequality, we can derive the evidence lower bound (ELBO): log *p*(*y*|**X, Z**, Θ_*n*_, Θ_*k*_, Θ_*p*_) *≥* E_*q*(**f**)_ log *p*(*y*|**f**) *−* KL *q*(**u**)||*p*(**u**) . The likelihood term is intractable. To address this, we first compute the marginal variational posterior distribution of **f**, denoted as *q*(**f**) = *N*(*µ*_*f*_ , Σ _*f*_ ). Then, we apply the reparameterization trick^72^ to *q*(**f**). We define **f** = *v*(*ε*_*f*_ ) = *µ*_*f*_ + **L** _*f*_ *ε* _*f*_ , with *ε* _*f*_ *∼ N*(**0, I**) and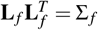. With this reparameterization, we can sample from the standard Gaussian distribution and approximate the likelihood term with Monte Carlo (MC) method^73^: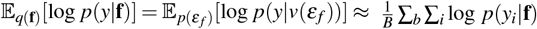, where *B* is the number of MC samples. With the above approximations, our model parameters can be efficiently learned by maximizing the ELBO using a first-order optimization method.

#### Post-hoc explanation

We design a novel technique for global model interpretation using the trained model. Our method can identify the most influential features/markers contributing to the final prediction. The identification of these markers could provide a more comprehensive understanding of the biological mechanisms underlying the clinical outcomes. More specifically, our explanation goal is to search for a minimal set of markers that effectively maintain prediction accuracy across all subjects. We consider these markers to be globally significant since, regardless of perturbations applied to the remaining markers, as long as the values of the identified markers are preserved, the model’s prediction remains largely unchanged.

This procedure can be formulated as an optimization problem, where the objective is to discover the minimum subset of markers that ensure high prediction performance when perturbing the values of the remaining markers. Formally, we use a mask matrix **M** *∈* ℝ^*M×p*^ to indicate marker importance. For the identification of globally important markers, we constrain each column of **M** to be the same value, **M**_:, *j*_ = 1 indicating the *j*-th marker is important and its value needs to be preserved, and **M**_:, *j*_ = 0 otherwise. The optimization problem can be formulated as finding the optimal **M** by solving the objective function: 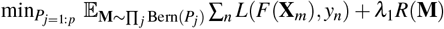,where *F* denotes our proposed model. We assume the value of each column **M**_:, *j*_ is independently sampled from a Bernoulli distribution parameterized by *P*_*j*_ (Bern(*P*_*j*_)). This design guarantees the value of **M**_:, *j*_ to be either 0 or 1 and enables the stochastic searching for the proper mask. The joint distribution is ∏ _*j*_ Bern(*P*_*j*_), where the entire matrix **M** is sampled from.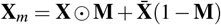, where **X** *∈* ℝ^*M×p*^ is the concatenation of original input cells of all samples across all time points. **X**_*m*_ is a perturbation of **X**, where the values of non-important markers are replaced with the mean value across all cells. Thus, the first term in the loss function *L* aims to minimize the prediction loss of the perturbed sample. For the second term, *R*(**M**) is the lasso regularization that restricts the number of non-zero elements in **M** and *λ* is a hyperparameter that controls the strength of *R*(**M**). The objective function can be hard to optimize because we cannot sample from the Bernoulli distribution with unknown parameters, and the Bernoulli distribution is discrete. To tackle these challenges, we approximate the Bernoulli distribution with its continuous relaxation – Gumbel-Softmax distribution^72^, which samples *e* _*j*_ from a uniform distribution *U* (0, 1) and computes **M**_:, *j*_ as a function of *P*_*j*_ and *e* _*j*_. With this approximation, we can move the unknown parameter *P*_*j*_ inside the expectation and sample from a known distribution *U* (0, 1) to solve *P*_*j*_ with a first-order optimization method. We use *P*_*j*_ as the final marker importance score, i.e., a larger *P*_*j*_ indicates the *j*th marker is more important.

#### Model parameters and tuning

In the cytoGPNet model, we begin by processing input cells across all subjects using an AE with a single hidden layer of 16 neurons, generating a latent embedding of dimension four. Our GP layer includes *O* = 100 inducing points and utilizes the SE kernel function, with the output variance set to 1 and the length scale set to 0.1. For AE pretraining, we set the batch size as 128, the learning rate as 1*e −* 6, the training epoch as 1000, and the optimizer as Adam. During end-to-end fine-tuning, we adjust the learning rate to 1*e −* 4, reduce the batch size to 10, and train for 100 epochs on the cytometry datasets. All the hyper-parameters are obtained through a grid search strategy. For scRNA-seq data, we modify the AE architecture by adding three fully connected hidden layers with output sizes of 256, 64, and 16, using the ReLU activation function. The widths of these hidden layers are pre-specified and not tuned. Through a grid search strategy, we set the batch size to 256, the learning rate to 1*e −* 3, and the training epoch to 200, using the Adam optimizer for AE pertaining. During end-to-end fine-tuning, we adjust the learning rate to 1*e−* 4, reduce the batch size to 10, and train for 100 epochs.

### Datasets

#### SDY1708

A total of 64 patients were enrolled in the Stanford University COVID-19 Biobanking studies from March to June 2020. All patients were over 18 years old and had a positive SARS-CoV-2 test result from the RT-PCR of a nasopharyngeal swab. Additionally, 8 asymptomatic adult donors were included as healthy controls in this study. Thus, the total number of individuals in the dataset is *N* =72, where 64 are COVID-19 positive and 8 are healthy controls. For all 72 individuals, blood was collected into cell preparation tubes or heparin vacutainers (*T* =1), and the CyTOF processing of 49 marker panels (*p* =49) was performed on peripheral blood immune cells preserved in liquid nitrogen to generate the single-cell mass cytometry data^74^. In our analysis, we let *y*_*n*_ =1 represent COVID-19 positive and *y*_*n*_ =0 represent healthy control.

#### SDY212

This dataset contains *N* =76 subjects who were enrolled in the influenza vaccine study. Peripheral blood samples, both pre- and post-vaccine (*T* =2), were collected. Flow cytometry analysis of eight marker panels (*p* =8) from experiment of EXP13405. Based on seroconversion, 52 individuals were classified as poor responders (PR), and the remaining were classified as good responders (GR)^75^. In our analysis, we let *y*_*n*_ =1 represent GR and *y*_*n*_ =0 represent PR.

#### HEUvsUE

This dataset contains *N* =44 African infants, comprising 20 who were either exposed to HIV in the maternal body but remained uninfected (HEU) and 24 who were unexposed (UE)^76^. A total of 308 blood samples were collected from these infants at six months after birth and were either left unstimulated (serving as a control) or stimulated with six Toll-like receptor ligands. Flow cytometry data with eight-marker panels (*p* =8) were obtained from all 308 blood samples. In our analyses, we treated *N* =44 using the unstimulated blood samples, and *N* =308 by aggregating all blood samples across the seven conditions. We defined *y*_*n*_ =1 for HEU infants and *y*_*n*_ =0 for UE infants.

#### TOP1501

*N* =29 patients with stage 1B-3A non-small cell lung cancer (NSCLC) received 2 cycles pembrolizumab, surgery, adjuvant chemotherapy, and 4 cycles of pembrolizumab on phase II study (NCT02818920). Viable and functional PBMCs were collected and stored at baseline and after pembrolizumab (*T* =2). Major pathologic response was observed in 7 of 29 patients after pembrolizumab. Functional immune cells were measured by flow cytometry using 25 marker panels (*p* =25) at baseline and after pembrolizumab. In our analysis, we use T cells identified by manual gating as input and let *y*_*n*_ =1 represent the major pathologic response (responder), and *y*_*n*_ =0 represent the non-major pathologic response (non-responder).

#### CMV

This longitudinal dataset comprises PBMC CyTOF files from an NK-centric panel (singlet or live cells post-debarcoding), collected from 11 CMV-viremic patients and 9 non-viremic (NV) patients^68^, for a total of *N* =20. Each NV patient has 3 FCS files corresponding to three distinct time points of blood sample collection. Each CMV-viremic patient has 4 to 5 FCS files representing five sampling time points, with four patients missing one collection. Due to sampling discrepancies between CMV-viremic and NV patients, and the absence of intermediate samples for certain CMV-viremic patients, our analysis focused on three standardized time points: day 0, pre-viremia, and post-viremia, to facilitate comparative analysis. Mass cytometry data with a total of *p* =39 marker panels were generated from these participants. We define *y*_*n*_ =1 to indicate a CMV-viremic patient and *y*_*n*_ =0 to denote a NV control subject.

#### SC4

SC4 is a large COVID single-cell RNA sequencing (scRNA-seq) dataset comprising *N* =196 subjects: 25 healthy controls and 171 COVID patients. Among the COVID patients, 79 exhibited mild or moderate symptoms, 92 were hospitalized with severe symptoms^77^. The dataset includes scRNA-seq data from 284 PBMC samples, processed using the 10x Genomics 5’ sequencing platform. In total, 27,647 genes were detected, from which 1,000 highly variable genes were selected for our subsequent analyses. In our analysis, we define the binary outcome variable *y*_*n*_ such that *y*_*n*_ =1 indicates a COVID positive patient, and *y*_*n*_ =0 indicates a healthy control.

#### Data preprocessing

To preprocess the cytometry data, we utilize the hyperbolic arcsine (arcsinh) transformation. This transformation is advantageous because it behaves similarly to a logarithmic transformation for high values while maintaining linearity around zero. Unlike the logarithmic transformation, arcsinh can effectively handle zero and small negative values. A key adjustable parameter of the arcsinh transformation is known as the ‘cofactor’, which controls the width of the linear region around zero. The transformation is defined as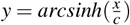, where x is the raw measured marker intensity value, and *c* is the cofactor. It is recommended to use a cofactor of 5 for CyTOF, and 150 for flow cytometry^78^. To preprocess scRNA-seq data, we perform several steps to ensure data quality and prepare for downstream analysis. First, we filter out genes that are expressed in fewer than 10 cells. Next, we normalize the data to account for differences in sequencing depth across cells, scaling the counts in each cell to a total of 10,000. We then apply a logarithmic transformation(`log1p`) to stabilize variance and approximate normal distribution of the expression values. The `log1p` transformation computes log(1 + *x*) for each expression value *x*. Finally, we identify the top 1,000 highly variable genes for our analysis^79–81^.

#### Comparison methods

#### CellCNN

CellCNN is an end-to-end model designed to identify rare cell subsets associated with potential disease status within a large number of cells^31^. The model employs a representation learning approach, enabling automatic feature extraction and deep learning simultaneously. The filter layers within the convolutional neural network (CNN) extract significant subject-level phenotypes from raw cytometry data inputs. To ensure that each sample contains an equal number of cells, CellCNN performs random subsampling from the original dataset before each run.

However, compared with later works in cytometry data modeling and analysis, CellCNN only captures limited cellular complexity^32^ and does not generalize well to datasets with more significant batch effects and measurement noises^33^. Moreover, the model supports only basic interpretability by identifying cell populations that most frequently activate each convolutional layer over many runs. CellCNN also does not utilize any form of temporal cytometry data in its predictions.

We downloaded Python codes for CellCNN from its GitHub repository: https://github.com/eiriniar/CellCnn/tree/python3. All four preprocessed datasets in CSV format were fed into the CellCNN model. We performed extensive hyperparameter tuning by exploring different combinations of key parameters: maxpool percent *∈* {0.01, 1, 5, 20} , number of filters *∈* {4, 6, 8, 10 }, learning rate *∈* {1*e −*4, 5*e −*4, 1*e −*3*}* , dropout probability *∈ {*0.1, 0.25, 0.5*}* , and L2 regularization coefficient *∈* {1*e−* 4, 1*e −*3, 1*e −*2} . For the TOP1501 dataset, which contains temporal information, we combined the cytometry data across different time points for each individual patient into a single data file, as CellCNN is unable to handle time-related information of cells.

#### CyTOF-DL

CyTOF-DL is an end-to-end model that uses a deep convolutional neural network to predict subject-level outcomes from cytometry data^32^. This model architecture captures the high-dimensional characteristics of cells through hidden layers and achieves invariance to cell permutation by using single-cell filters and max/mean pooling. The model also requires subsampling to keep the number of cells equal within each sample. To extract useful biological insights from the trained model, a permutation-based interpretation pipeline was developed. This pipeline quantifies how much each cell in the dataset influences the model’s prediction outcome. However, the model does not support the input of temporal cell information and exhibits a significant drop in prediction accuracy when the sample size is less than 200 or when fewer than 8 cell markers are used^32^.

We downloaded Python codes for CyTOF-DL from its GitHub repository: https://github.com/hzc363/DeepLearningCyTOF. All four preprocessed datasets in CSV format were fed into the CyTOF-DL model. During model optimization, we experimented with different architectural configurations: convolutional filters *∈* {3, 8, 16 }, dense units *∈* {3, 16, 32 }, and learning rates *∈ {*1*e −*4, 5*e −*4, 1*e −*3*}* . For the TOP1501 dataset, which contains temporal information, we combined the cytometry data across different time points for each individual patient into a single data file, as CyTOF-DL is unable to handle time-related information of cells.

#### CytoSet

CytoSet is an end-to-end model designed for clinical outcome prediction, utilizing a custom permutation-invariant architecture^33^. CytoSet treats a collection of cells in a dataset as an unordered set, rather than a regular ordered matrix, based on the assumption that the order in which cells are profiled does not have significant biological implications. Due to this unordered approach to dataset handling, the CytoSet architecture does not consider temporal information. CytoSet also requires an equal cell count across different samples, necessitating subsampling before training. Additionally, the model lacks interpretability features.

We downloaded Python codes for CytoSet from its GitHub repository: https://github.com/CompCy-lab/cytoset. Similar to the above methods, all four preprocessed datasets in CSV format were fed into the CytoSet model. We conducted hyperparameter tuning by varying the number of blocks *∈* { 2, 3, 4 }, hidden dimension sizes *∈* { 128, 256, 512 }, learning rates *∈* { 1*e −*5, 5*e −*5, 1*e −*4} , *β*_1_ *∈* { 0.85, 0.9, 0.95 }, *β*_2_ *∈* {0.99, 0.999, 0.9999} . The parameters *β*_1_ and *β*_2_ are the exponential decay rates for the moment estimates in the Adam optimizer. *β*_1_ controls the decay rate of the first moment (mean) of the gradient, while *β*_2_ controls the decay rate of the second moment (uncentered variance) of the gradient. These hyperparameters affect how quickly the optimizer adapts to changes in the gradient during training. For the TOP1501 dataset, which contains temporal information, we combined the cytometry data across different time points for each individual patient into a single data file, as CytoSet is unable to handle time-related information of cells.

#### AE

We design this AE model as a baseline for comparison with cytoGPNet. Like cytoGPNet, we use AE to encode the cytometry data. However, unlike cytoGPNet, we bypass the GP layer and instead pass the latent output directly to the set of attention layers and logistic regression to generate subject-level outputs. Similar to cytoGPNet, the AE is first pre-trained by applying reconstruction loss to the cytometry data at the single-cell level with a default learning rate of 10^*−*6^ under 100 epochs. Subsequently, the parameters of the AE are fine-tuned along with the remaining layers by minimizing the binary cross-entropy loss based on the subject-level outputs. To process temporal data, all cells will be collectively passed through a single AE. Subsequently, they will be directed to distinct attention layers corresponding to each time point. These time-specific representations will then serve as separate covariates for logistic regression in the generation of subject-level predictions.

#### Logistic regression

Logistic regression (LR) is a widely utilized parametric model for analyzing multivariate data (features) with binary outcomes, as is the case in our paper. LR assumes a linear relationship between the logit transform of the class probability and the covariates. In our analysis, we employed mean expression values for all the markers of each subject as features to predict subject-level outcomes. For the cytometry data collected across time, we concatenated the mean expression values of all markers at each time point to form the full set of features. We used the `glm` function in R to implement LR and did not apply any regularization in our analysis.

#### Random Forest

Random forest (RF) is an ensemble classification method that utilizes decision trees as its baseline classifiers. Each decision tree is trained on a random sample with replacement of the original data. Moreover, during the split of each node in a decision tree, only a random subset of features is considered. A key advantage of RF is its resistance to overfitting, achieved through the voting mechanism on decisions from multiple trees. Similar to the use of the above LR, we used mean expression values for all the markers as the feature set and applied the `randomForest` package in R with the default settings of 500 trees and randomly sampled 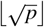 features at each split, where *p* is the total number of features.

## Conclusion

In this study, we introduce cytoGPNet, a novel approach designed to enhance the prediction accuracy of individual-level clinical outcomes using longitudinal cytometry data, particularly under the constraints of limited sample sizes. Traditional cell-gating techniques, while prevalent for extracting salient features from cytometry data, often lack direct integration with predictions, thereby reducing the relevance of cell-gating outcomes to the desired clinical results. In contrast, deep learning models leverage the extensive cytometry data directly, employing back-propagation algorithms to iteratively adjust parameters, thus refining feature detection pertinent to the desired outputs. Deep learning models show significant potential for analyzing cytometry data. However, recent methods, despite their innovation, encounter challenges in achieving consistent performance across diverse cytometry datasets. This issue is especially pronounced in datasets with very few subjects, such as the TOP1501 dataset, where stability of performance metrics like AUC and F1 score can be compromised. Moreover, many existing methods require uniform cell counts per sample input, necessitating random sampling during preprocessing, which can result in loss of crucial cell information and reduce their effectiveness in handling longitudinal cytometry data.

Our developed method, cytoGPNet, addresses these multiple challenges inherent in cytometry data analysis: accommodating varying cell counts per sample, modeling temporal relationships, maintaining robustness with limited samples, and ensuring interpretability. cytoGPNet demonstrates substantial improvements over existing methods. Our application of cytoGPNet across diverse studies highlights its consistent superiority in prediction accuracy and its ability to provide valuable, interpretable insights. These findings underscore the potential of cytoGPNet to significantly advance the analysis of cytometry data, offering a powerful tool for immunological research. In addition to the strengths of our model, it’s important to acknowledge potential extensions and limitations. Incorporating demographic information as additional covariates in the regression layer, as is common in many DNN models, could further enhance its predictive power. We opted not to include this feature in our current model due to inconsistencies across the four datasets, with two lacking demographic data. However, any future design incorporating demographic variables should be carefully implemented in subject-level outcome prediction, particularly given the relatively small sample size, to avoid overfitting. Our current approach employs a unified GP to capture cell-cell correlations across different time points. We visualized the covariance matrix and cannot find clear patterns for cell correlations and dependencies. An extension of the current explanation mechanism could involve designing more detailed analysis methods that leverage the covariance matrix to uncover hidden patterns. For example, we can apply clustering methods and then find the patterns across different clusters. Meanwhile, our model significantly outperform in longitudinal data but may not improve that much on cytometry data with single time point. Our cytoGPNet framework’s flexibility is evident in its adaptability to different data types. For lower-dimensional cytometry data, the AE model effectively extracts cell representations. However, for higher-dimensional single-cell data, such as scRNA-seq, AEs may lose information. In such cases, transformer-based architectures could serve as alternative approaches for representation learning. Transformers have shown promise in modeling complex dependencies in high-dimensional data^82^. Nonetheless, transformers do not inherently perform dimensionality reduction, so additional steps would be necessary to integrate their outputs into the GP component.

## Data availability

The data that support this paper’s findings are available at ImmPort (https://www.immport.org) under study accession SDY1708 and SDY212, at FlowRepository (http://flowrepository.org) with repository ID FR-FCM-ZZZU for HEUvsUE, at Mendeley Data, (https://data.mendeley.com/datasets/fnbvcyf223/1) for CMV and at CELLxGENE (https://cellxgene.cziscience.com/collections/0a839c4b-10d0-4d64-9272-684c49a2c8ba) for SC4. Raw flow cytometry data for TOP1051 is publicly available on Zenodo at https://zenodo.org/records/15002576.

## Code availability

An open-source implementation of cytoGPNet is available on GitHub at https://github.com/llin-lab/cytoGPNet.

## Acknowledgements

Merck Sharp & Dohme LLC, a subsidiary of Merck & Co., Inc., Rahway, NJ, USA provided financial support for the study. The opinions expressed in this paper are those of the authors and do not necessarily represent those of Merck Sharp & Dohme LLC. This research was also supported by the Duke University Center for AIDS Research (CFAR), an NIH funded program (5P30 AI064518), and NIH P01 (2 P01 AI129859). The authors gratefully recognize the contributions of Jennifer Enzor and Prekshaben Patel, who generated all of the original TOP1501 flow cytometry data in the Duke Immune Profiling Core (DIPC), a designated Shared Resource of the NIH-sponsored Duke Cancer Institute (5P30-CA014236-50). We would like to thank Xiaoyu Liu for his valuable assistance in performing the benchmarking analysis, which contributed to the early stages of this project.

## Author contributions statement

JZ programmed the model, performed data analyses, and manuscript editing. LS performed benchmark analysis and manuscript editing. NR reviewed data and manuscript editing. WG conceptualized the new model and manuscript writing. LL conceived the study, conceptualized the new model, interpreted the results, and manuscript writing.

## Footnotes

1 We follow the common setup^70^ and set the mean function of GP as 0.

