## Supplementary for "cytoGPNet: Enhancing Clinical Outcome Prediction Accuracy Using Longitudinal Cytometry Data in Small Cohort Studies"

### Supplementary Information

| Dataset | Data Type | # Subjects | # Cells | Median # Cells per Subject | # Markers | Phenotype | # Time Points |
|---|---|---|---|---|---|---|---|
| **SDY1708** | CyTOF | 72 | 2,089,457 | 20,646 | 49 | COVID vs. Healthy | 1 |
| **SDY212** | Flow | 76 | 2,051,080 | 22,922 | 8 | Good responder vs. Poor responder to influenza vaccine | 2 |
| **HEUvsUE** | Flow | 308 | 135,638,183 | 460,161 | 8 | Exposed to HIV vs. Non-exposed to HIV | 1 |
| **TOP1501** | Flow | 29 | 1,164,636 | 41,547 | 25 | Major pathologic response (responder) vs. Non-major pathologic response (non-responder). | 2 |
| **CMV** | CyTOF | 20 | 6,595,269 | 315,524 | 39 | CMV viremic vs. Non-viremic | 3 |
| **SC4** | scRNA-seq | 196 | 1,462,702 | 5,939 | 27,647 | Mild or moderate vs. Severe COVID symptoms vs. Healthy | 1 |

**Table S1.** Summary information on the six datasets. For each dataset, data type, number of subjects, total number of cells, median number of cells per subject, number of markers, phenotype, and number of time points are listed.

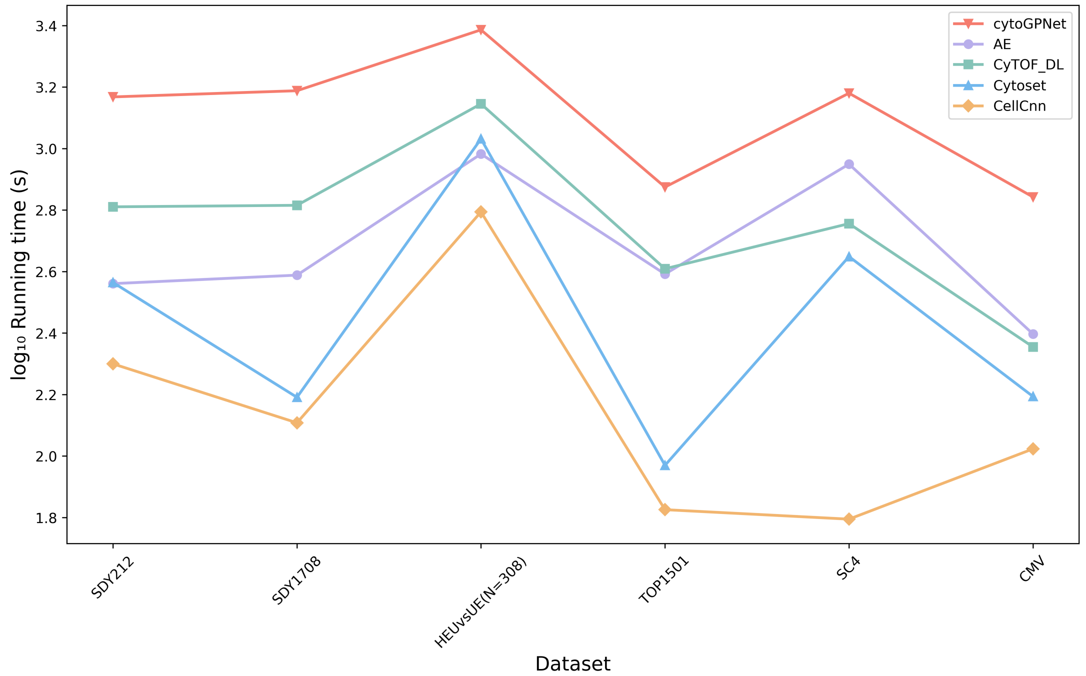


**Figure S1.** Comparison of computational time for cytoGPNet versus other deep learning-based methods across six datasets. cytoGPNet can be efficiently run on a 32GB GPU (NVIDIA RTX 5000 Ada Generation) paired with an 8-core Intel Xeon Gold 6336Y CPU @ 2.40GHz.

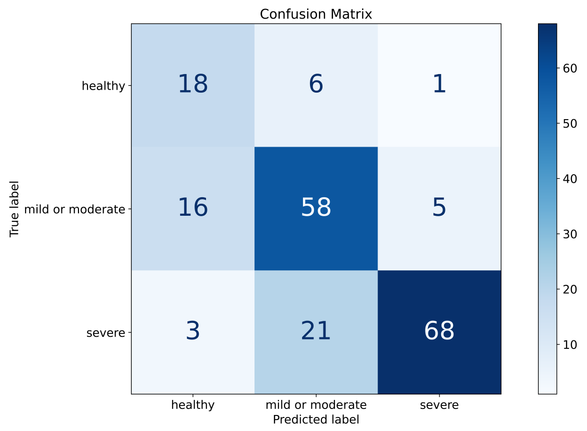


**Figure S2.** Confusion matrix for the cytoGPNet's classification of three classes for SC4 dataset: healthy, mild or moderate, and severe. The rows represent the actual class labels, while the columns indicate the predicted labels. The diagonal entries correspond to correctly classified instances, whereas off-diagonal entries indicate misclassifications. The color intensity and numerical values reflect the number of instances in each category, highlighting the model's performance across different classes.

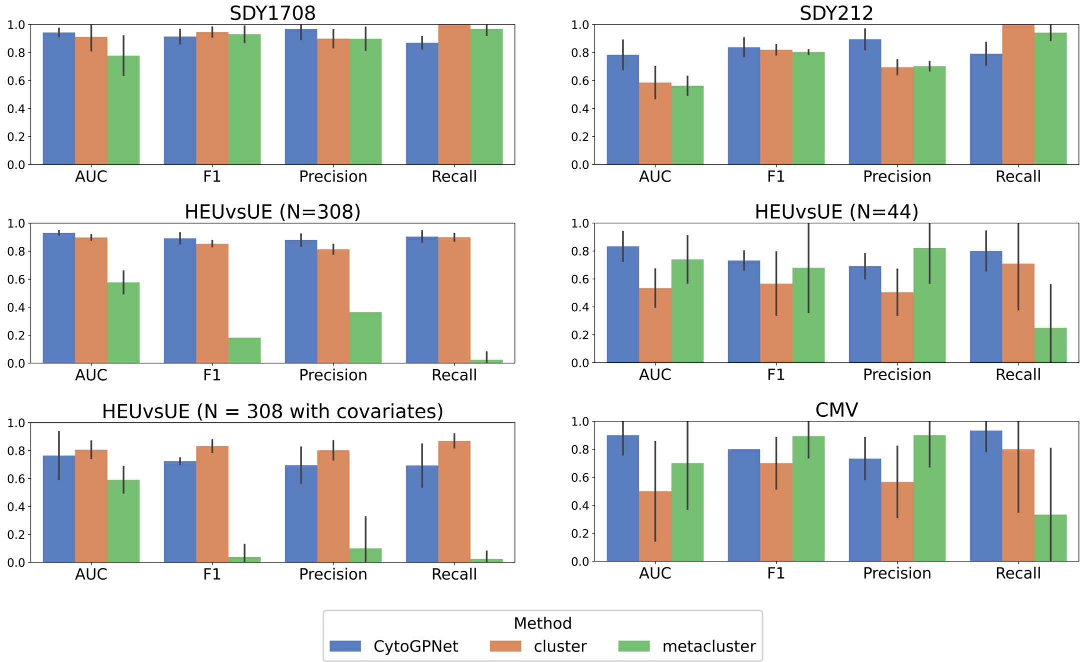


**Figure S3.** Comparison of cytoGPNet and LR with Lasso penalty using cell type proportions (clusters or meta-clusters) as features. Performance is evaluated using AUC, F1-score, precision, and recall based on 5-fold cross-validation across four datasets: SDY1708, SDY212, HEUvsUE, and CMV. The height of each bar represents the corresponding mean value across the five folds, and the vertical lines (error bars) indicate the standard error.

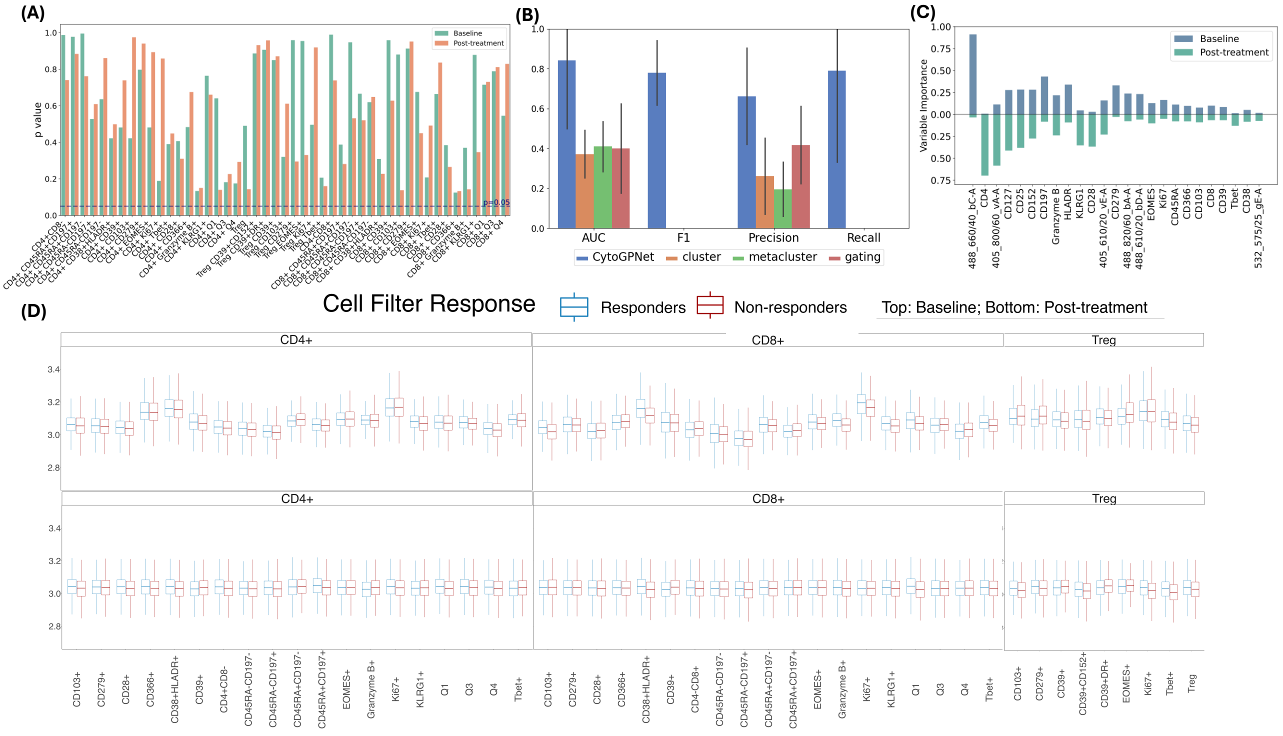


**Figure S4.** (A) Bar plots displaying p values from univariate logistic regression analyses, where each cell type proportion is added as a covariate. (B) Prediction performance of cytoGPNet and logistic regression with Lasso penalty based on cell type proportions obtained from FlowSOM (using both 100 clusters and 10 metaclusters) as well as manual gating results using 5-fold cross-validation. (C) Barplots representing variable importance for RF for both baseline and post-treatment data. (D) Boxplots comparing cell filter response values between responders and non-responders across all cell subsets in the CellCnn model.

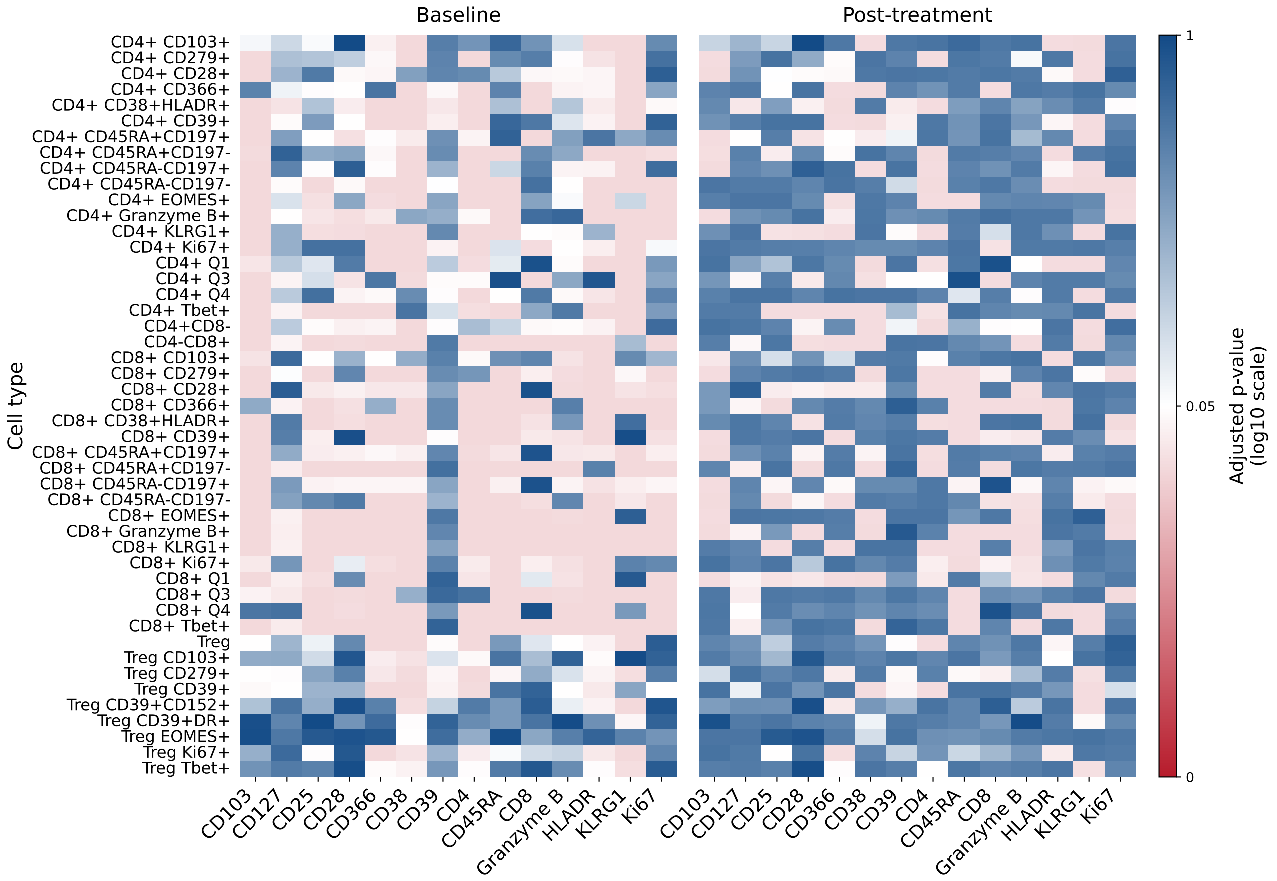


**Figure S5.** Heatmap visualization of adjusted p values from differential abundance analysis across manually gated cell types and markers in baseline and post-treatment conditions. p values were transformed using log10 scale to enhance visualization of significance levels. Values were generated using the diffcyt method, with statistical significance indicated by a color gradient: deep red (highly significant, p value near 0), white (p=0.05 threshold), and deep blue (non-significant, p value near 1).

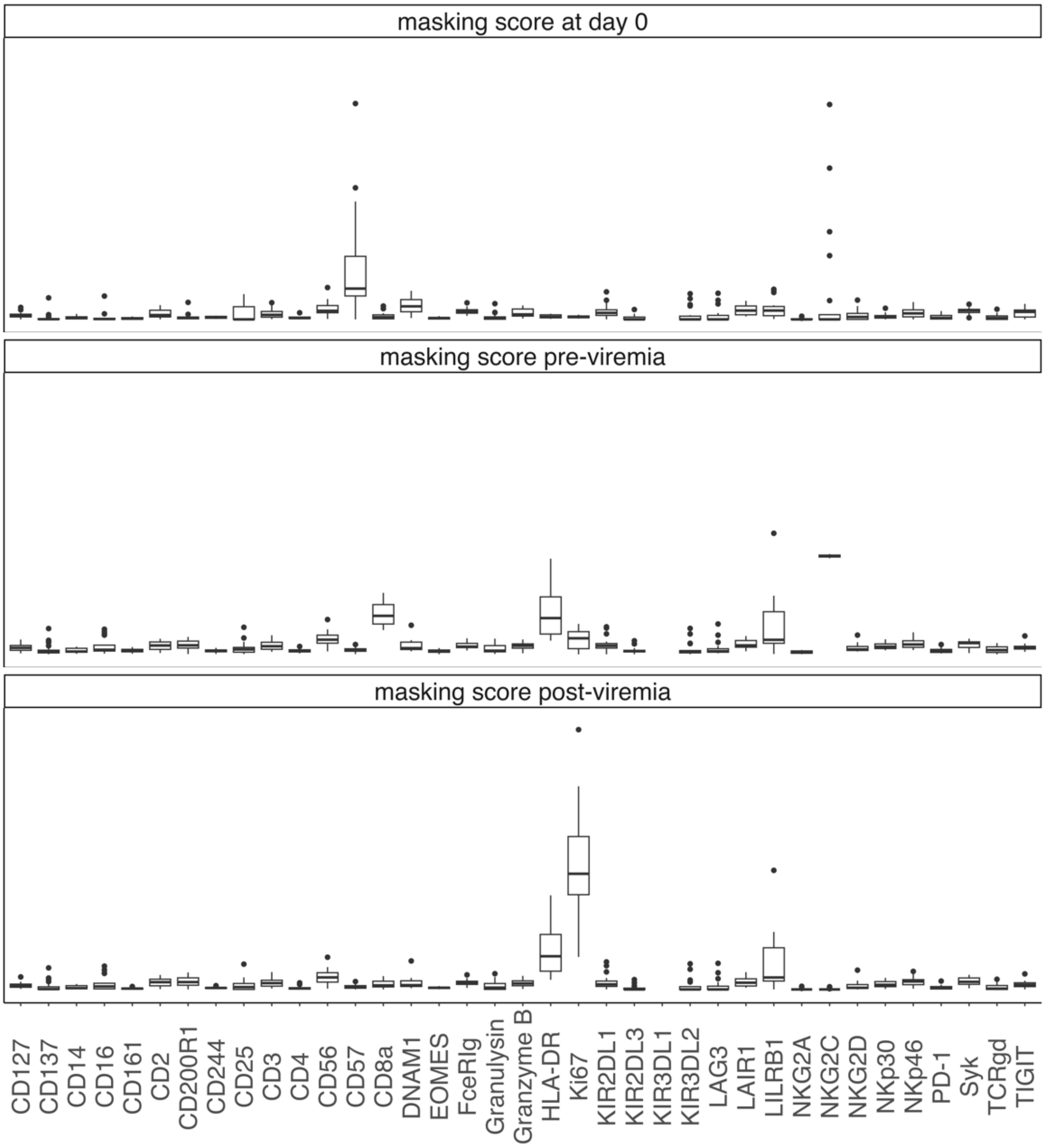


**Figure S6.** Boxplots visualizing the distribution of masking scores (ranging from 0 to 1) for each marker (x-axis) for CMV data at day 0 (top), pre-viremia (middle), and post-viremia (bottom).
